# Extracting robust trends in species’ distributions from unstructured opportunistic data: a comparison of methods

**DOI:** 10.1101/006999

**Authors:** Nick J.B. Isaac, Arco J. van Strien, Tom A. August, Marnix P. de Zeeuw, David B. Roy

## Abstract

1. Policy-makers increasingly demand robust measures of biodiversity change over short time periods. Long-term monitoring schemes provide high-quality data, often on an annual basis, but are taxonomically and geographically restricted. By contrast, opportunistic biological records are relatively unstructured but vast in quantity. Recently, these data have been applied to increasingly elaborate science and policy questions, using a range of methods. At present we lack a firm understanding of which methods, if any, are capable of delivering unbiased trend estimates on policy-relevant timescales.
2. We identified a set of candidate methods that employ either selection criteria or correction factors to deal with variation in recorder activity. We designed a computer simulation to compare the statistical properties of these methods under a suite of realistic data collection scenarios. We measured the Type 1 error rates of each method-scenario combination, as well as the power to detect genuine trends.
3. We found that simple methods produce biased trend estimates, and/or had low power. Most methods are robust to variation in sampling effort, but biases in spatial coverage, sampling effort per visit, and detectability, as well as turnover in community composition all induced some methods to fail. No method was robust to all forms of variation in recorder activity.
4. We warn against the use of simple methods. We identify three methods with complementary strengths and weaknesses that are useful for estimating timely trends. Sophisticated correction factor methods, including Occupancy and Frescalo, offer the greatest potential in the long-term. Methods based solely on selection criteria are inherently limited, but a combination or ensemble of approaches may be required to generate trends that are both robust and powerful. Small amounts of information about sampling intensity, captured at the point of data collection, would greatly enhance the utility of opportunistic data and make future trend estimates more reliable.

## Introduction

Robust quantitative measures of the stock and rate of change in biodiversity are crucial for assessing species’ risk of extinction (Mace & Lande, 1991), for measuring progress against international targets (Butchart et al., 2010) and testing against predictions about climate change impacts (Maclean & Wilson, 2011). The demands for timely information are increasing. For instance, the EU Habitat and Bird directives require changes in species’ status to be reported every six years, and progress against the Convention of Biological Diversity targets are reported on a decadal basis.

Long-term, standardized, monitoring schemes produce timely and robust estimates of status and trends, often on an annual basis (Gregory et al., 2005). Unfortunately such data are available for only a small number of taxa in a few countries. The next best sources are opportunistic data, such as those available on the Global Biodiversity Information Forum (GB1F), including records submitted by volunteers (Prendergast et al., 1993). These data are less structured than monitoring schemes but high in quantity: GB1F comprises 417 million observations of 1.4 million species (http://www.gbif.org). Opportunistic data have delivered substantive insights into the ecological impacts of climate change (Hickling et al., 2006), invasive species (Roy et al., 2012) and habitat loss (Warren et al., 2001).

Whilst opportunistic data have been used to describe coarse-scale changes in biodiversity (e.g. Carvalheiro et al., 2013; Thomas et al., 2004), the absence of standardized protocols presents serious challenges for estimating timely trends in the status of individual species. The noise generated by opportunistic sampling has the potential to swamp any signal of real change, or to produce spurious signals of change where none exists. We use the term ‘variation in recorder activity’ to refer to the sampling biases inherent in opportunistic data, of which there are four principle forms: 1) uneven recording intensity over time, measured as the number of visits per year (a visit is defined as unique combination of site and date in the records data), 2) uneven spatial coverage, 3) uneven sampling effort per visit, and 4) uneven detectability. Each source of variation has the potential to introduce substantial bias in trend estimates for individual species. The growth of citizen science programs (Dickinson et al., 2012) is likely to increase data volumes, and affect the nature of recording with potentially far-reaching consequences for how the data may be used to infer biodiversity trends (Tulloch et al., 2013).

In the past, opportunistic data were often treated by collating many years’ data in one Atlas period. This compensates to some degree for variation in recorder activity, allowing changes in species distributions to be assessed over the years between atlas periods (Botts et al., 2012; Thomas et al., 2004; Tingley & Beissinger, 2009). This approach has limited potential to deliver trends in a timely fashion, because Atlas periods are typically measured in decades. In principle, it should be possible to derive trend estimates on sub-decadal timescales by incorporating information about the data collection process (Roy et al., 2012; Szabo et al., 2010; van Strien et al., 2013). Therefore, a pressing need exists to understand how recorder activity can be treated statistically. Identifying methods that are robust would open a vast frontier of previously unexploited data for use in both biodiversity policy and applied ecology.

There are numerous methods proposed in the literature for estimating trends in species’ distributions from opportunistic data whilst taking into account recorder activity. Here we test a representative set of methods under realistic scenarios of recorder activity. Our aim is to identify methods that produce timely trends that are robust to multiple forms of variation in recorder activity.

## Range change methods

Many metrics have been proposed to account for variation in recorder activity when estimating trends in species’ distributions from opportunistic data (table 1). Methods also differ in the spatial and temporal resolution at which they are applied, but we focus on the underlying assumptions they make. Technical details of all the methods, including mathematical notation, can be found in Appendix S1.

**Table 1:** Methods to estimating trends in distribution in opportunistic data and the way they control for variation in recorder activity.

| Category | Source | Name | Metric of change | Temporal scale | Mechanism to adjust for variation in recorder activity |
| --- | --- | --- | --- | --- | --- |
|  | This study | Naïve | Number of sites | year | None |
| Selection | Maes et al. (2012) | RDC index | Proportion of unique records on well-sampled sites | period | selection of well-surveyed sites + surveyed in each period + taking into account sum of sites for all species |
|  | Roy et al. (2012) | Well-sampled Sites | Probability of being recorded per visit | year | 'Well-sampled sites' defined by threshold list length per visit and number of years visited + site effect |
| Correction | Ball et al. (2011) | Reporting Rate | Probability of being recorded per visit | year | Expressing the records as a proportion controls for temporal variation in number of visits |
|  | Szabo et al. (2010) | List Length | Probability of being recorded per visit | year | Number of species per list (the list length) as proxy for sampling effort of each visit |
|  | Telfer et al. (2002) | Telfer index | Number of sites | period | Difference in the number of sites per period is expressed relative to that across other species |
|  | Hill (2012) | Frescalo | Relative reporting rate | period | Detection of regional benchmark species as proxy for recorder effort |
|  | Van Strien et al. (2013) | Occupancy | Probability of occupancy | year | Detection probability as proxy for recorder effort |

The simplest measure of change is the linear trend (or difference) in the annual number of sites (or grid cells) occupied by the species of interest (i.e. a Poisson generalised linear model (GLM)). This model has no mechanism to control for recorder activity, so we refer to it as the *Naïve* method. The *Naïve* method is unique in that the trend is based solely on records from the focal species. All others employ records from other species to control for variation in recorder activity, either assuming that a record of one species indicates the absence of others, or as a means of estimating sampling effort.

The methods available to cope with variation in recorder activity fall into two broad categories: employing selection criteria or applying correction factors (table 1). The rationale behind selection criteria is that it is possible to select a subset of records that are free from bias (Botts et al., 2012). Many selection methods have been proposed (Hickling et al., 2006; Kuussaari et al., 2007; Maes & Van Dyck, 2001; Maes et al., 2012; Rich & Woodruff, 1996; Roy et al., 2012; Van Calster et al., 2008; Warren et al., 2001): we chose two representatives for closer examination.

Maes et al. (2012) applied the criterion that grid cells should have at least five species recorded in each of two time periods. This provides a simple way to correct for both the number of visits and effort per visit. For each period the relative distribution for a species is the proportion of unique records (period-cell-species combinations). The *Relative Distribution Change* (*RDC*) index is the difference in relative distribution between the two time periods, divided by the value in the first time period.

Roy et al. (2012) used a mixed-effects model to explore the impacts of an invasive ladybird on native species. They defined thresholds of two species per visit and three years per site for including data within the ‘well-sampled’ subset. We use a modified version of this model, which we refer to as the *Well-Sampled Sites (WSS)* method (see Appendix S1 for details). The observations are unique combinations of site and year, with a binomial response variable for estimating a trend in the probability of being recorded on an average visit. We expect that WSS is likely to perform badly when sampling effort per visit changes over time. We test two versions, *WSS_2* and *WSS_4,* where 2 and 4 indicate the threshold number of species per visit to meet the well-sampled criterion.

The second category of methods has a statistical correction procedure to treat recorder activity. These methods are less frequent in the literature than selection methods, but have a greater variety of mechanisms to control for recorder activity. To cover this variety we selected five methods for comparison (table 1).

Ball et al. (2011) proposed a simple improvement to the *Naïve* model to control for changes in overall recording intensity over time. The *Reporting Rate* is the proportion of visits on which the focal species was recorded, under the assumption that the effort per visit does not vary among years. We implemented two variants: *ReportingRate* is a binomial GLM and *ReportingRate+Site* incorporates a random effect for site identity, which is equivalent to the *WSS* model without selection criteria. We predict that both variants are robust against variation in the number of visits, but will be sensitive to uneven sampling effort per visit. To address this problem, Szabo et al. (2010) proposed a modification in which individual visits (or species lists) are the unit of analysis (thereby controlling for variation in the number of lists over time). Their innovation was to treat the number of species on the list (the list length, L) as a proxy for recorder effort per visit. We use the GLM version of the *ListLength* method, as well as a *ListLength+Site* variant with random effect for site. We predict that *ListLength* will be robust to trends in both the number of visits and the sampling effort per visit.

Telfer et al. (2002) used the estimated trend in all species together as an indirect measure of how recording intensity differed between two sampling periods. If recorder intensity is higher in the second period, all species are expected to show increases compared with the first period. Any deviation from the overall expected trend is considered as an index of change for the species of interest. The *Telfer* index for each species is the standardised residual from a linear regression and is a measure of relative change only, because the average real trend across species is obscured. We predict that *Telfer* will be sensitive to scenarios in which recording is biased with respect to the focal species (e.g. spatial bias or changes in detectablity).

Both Maes & van Swaay (1997) and Hill (2012) developed methods using benchmark species as proxy for recorder activity. Benchmarks are common species whose distribution is assumed to show no overall trend. We selected Hill’s method, known as *Frescalo,* which uses information about sites’ similarity to one another to assign local benchmarks within neighbourhoods, and provides site-specific estimates of recording intensity. We compare two variants: in *Frescalo_P* we pooled the data into two equal time periods; in *Frescalo_Y* the data were analysed in ten time-periods (i.e. one per year). *Frescalo* trends are expressed as the reporting rate of focal species relative to that of the benchmarks (see Hill 2012 for further details). We predict the performance of *Frescalo* will be similar to *Telfer’s* method, but more powerful.

Finally, we included *Occupancy* modelling (MacKenzie, 2006) in our study. Occupancy models are derived from capture-recapture theory and have recently been successfully applied to large-scale models of distributional change (Van Strien et al., 2013). The key feature of Occupancy is that it uses replicated visits within a season to estimate the probability that a species is recorded when present. The model consists of two hierarchically coupled submodels, one governing occupancy (presence-absence) and the other governing the observations (detection-nondetection). Following van Strien et al. (2013), the observation submodel includes a covariate for sampling intensity per visit, based on the list length, L (see Appendix S1 for full details). This statistical separation of detection from presence-absence represents a major advance (MacKenzie, 2006) and we predict that *Occupancy* will be the most robust method to be tested. As with other methods, we tested a simple *Occupancy* model and an *Occupancy+Site* variant.

## Simulation design

We constructed a computer simulation to assess the performance of the proposed methods under simple deviations from a control scenario of random sampling. We generated species occurrence matrices using simple rules, which were then subjected to a suite of recording scenarios by virtual observers (Zurell et al., 2010) to generate a set of realised datasets. Our recording scenarios simulate temporal trends in recorder activity, as well as changes in community composition. Where possible, our scenarios were parameterised using observed patterns of recording in the Great Britain and the Netherlands (Isaac, 2012; van Strien et al., 2010). We then estimated a trend in the distribution of one ‘focal’ species on each realised dataset using the methods described above. The performance of each method-scenario combination was assessed from 500 simulated datasets. We conducted separate tests of each method’s validity and its power to detect change.

## Species occurrence data

Our system consists of 1000 ‘sites’, which we assert to be separated in space (although our simulation for simplicity’s sake is not spatially explicit). Each test dataset consisted of one focal species and 25 non-focal species (preliminary analyses showed the results are insensitive to the total number of species). Species were distributed randomly among sites: each distribution was determined by drawing 1000 times from a binomial distribution with a species-specific probability of being occupied. For the focal species’ we fixed this probability at 50% in all simulations; for non-focal species we used random numbers from a beta distribution with shape parameters 1 and 2, such that mean species richness among sites was ∼13 species, with a variance among sites of ∼5. We ran all simulations over a period of 10 years.

## Control scenario

This section defines the *Control* scenario, which corresponds to random sampling. Most departures from random sampling were generated by subsampling from the records generated by the *Control.*

Overall recording intensity was characterised by the number of visits each year. Within years, the distribution of visits among sites is characterised by a power law decay, i.e. some sites receive many visits and most sites receive few (with a mode of zero). In a selection of British and Dutch recording datasets, the power law exponent is close to −2, indicating that the number of sites receiving *n* visits is 4 times greater than number receiving 2*n* visits (2^−2^ = 0.25).

Variation in the total recording intensity is characterised by the proportion of sites that receive a single visit each year (i.e. the intercept in the power law function). We selected three levels of overall recording intensity (low, medium, high), corresponding 5%, 7% and 10% of sites that receive a single visit each year. This range of values was selected in order to generate datasets that superficially resemble the records of dragonflies (high intensity) and beetles (low intensity) in the UK.

Each year, a team of virtual observers visited a certain number of sites. Sites were selected by sampling a multinomial distribution defined by the power law function above, truncated so that no site received more than 10 visits in any one year. The number of sites to be visited varies from year to year, but the parameters of the power law were constant across years. Although sites were selected at random, the visits were apportioned non-randomly: specifically the number of visits to each site was determined by its species richness, with the most speciose site receiving most visits. This was done in order to mimic real datasets in which records are clustered around nature reserves and other sites that are known to harbour interesting wildlife.

Species do not automatically get recorded if a site is visited, since most surveys are incomplete (Isaac, 2012; van Strien et al., 2010) and many species are rarely encountered. Each species had a fixed probability of being detected if present (i.e. we assume that visits have equal sampling effort). The focal species detection probability was fixed at 0.5 per visit; for nonfocal species the detection probability varied from 0.88 - 0.16 following the sigmoid curve described in Hill (2011). This species-specific detection probability can be thought of as the product of visual apparency (Dennis et al., 2006) and mean abundance. Species’ detection probabilities were uncorrelated with occupancy.

Low and high recording intensity delivered 38 and 77 records per species per year, respectively, under this *Control* scenario. Other scenarios produce recording rates that are comparable with the four decade average of 20 records per species per year across a range of taxa regarded as moderately well-recorded in the UK (Isaac, 2012).

## Biased recording scenarios

We devised five biased recording scenarios (table 2) to capture the four major axes of variation in recorder activity, as well as changes in community composition.

**Table 2:** Description of recording scenarios in the simulation

| Scenario | Summary |
| --- | --- |
| *Control* | Constant recording intensity over years. All species have a fixed probability of being recorded per visit. |
| *MoreVisits* | Number of visits per year doubles over the course of the recording period. |
| *MoreVisits+Bias* | As *MoreVisits*, but the extra visits are biased toward sites where the focal species is absent. |
| *LessEffortPerVisit* | Sampling effort per visit declines over time, increasing the proportion of 'short lists' from 60% to 90% of visits. |
| *MoreDetectable* | The focal species is 20% easier to detect at the end of the recording period than at the start. |
| *NonFocalDeclines* | 50% of nonfocal species are each declining at 30% over the recording period. |

The first simulates an increase in the number of visits per year (i.e. recording intensity is uneven over time). In the *MoreVisits* scenario the expected number of visits per year doubled over the ten year recording period. We simulated this by sub-sampling from the *Control* scenario: each year we sampled (without replacement) a proportion of visits, with the proportion in the final year set equal to 1. Our second scenario, *MoreVisits+Bias,* is a modification in which sites are selected nonrandomly: this simulates temporal change in the spatial coverage of sites. Specifically, sites containing the focal species are 27% more likely to be visited (than non-focal sites) in year one, but in year 10 the focal and nonfocal sites are equally represented.

Uneven sampling per visit is the third major axis of variation in recorder activity. Inter-annual variation in sampling effort is a potentially serious form of bias for some methods, because it affects species’ probabilities of being recorded. We simulated a directional trend towards shorter lists, as might result from changes in recorder behaviour (e.g. a growth in the number of inexperienced recorders with limited identification skills). In the *LessEffortPerVisit* scenario, the prevalence of short lists increased from 60% to 90% in each simulation. Short lists contained 1, 2 or 3 species, in the ratios 2:1:1 respectively. As above, this was achieved by subsampling from data produced under the *Control* scenario. The total number of records produced by *LessEffortPerVisit* is around half the number produced by the *Control.*

We also model situations in which species become more detectable over time, e.g. through the adoption of new technology or publication of a field guide. In the *MoreDetectable* scenario, we model a gradual increase in the focal species’ probability of detection per visit, from 0.4 at the start of the simulation to 0.5 at the end (i.e. a 20% increase over the recording period).

Several of the methods described above measure relative, rather than absolute, change (*Telfer, ListLength* and *Frescalo*). For this reason, an important consideration is the degree to which these relative trends are impacted by changes in the status of other (nonfocal) species. We tested this by simulating a decline of 50% over ten years in 30% of nonfocal species (*NonFocalDeclines*). Declining species were selected at random in each simulation.

## Estimating the trends and evaluating model performance

For each simulated dataset we tested the null hypothesis of no change in the focal species’ distribution using each of the 13 method variants. Full details of how we derived p-values for each method are described in Appendix S1. For *RDC, Telfer* and *Frescalo_P* we split the realised data into two five-year periods. To implement *Frescalo* we generated a random matrix of neighbourhood weights: randomly-generated neighbourhoods would be inappropriate for real datasets where communities show strong evidence of species sorting, but are reasonable for our simulated data in which species were independently distributed. Other parameters of *Frescalo* were set following Hill (2012). We implemented *Occupancy* in a Bayesian framework using JAGS with three Markov chains, 5000 iterations per chain, a burn-in of 2500 and a thinning rate of three (van Strien et al., 2013).

For the test of validity, the distribution of the focal species remained unchanged throughout the simulation: the Type 1 error rate is the proportion of 500 simulated datasets in which the null hypothesis was rejected at α=0.05. In the test of power we simulated a linear decline in occupancy of the focal species of 30% over the 10 year period (i.e. the species would qualify as Vulnerable under IUCN Criterion A2). A simple estimate of power would be the rate at which we failed to reject the null hypothesis (the Type 11 error rate). However, some scenarios are designed to introduce negative bias in the trend estimates, so Type 11 error rates are not comparable across scenarios. Instead we defined power as the proportion of simulations in which a true decline was successfully detected (at α=0.05) minus the matching Type 1 error rate, with a lower boundary of zero.

## Results

About half the methods return appropriate Type 1 error rates (α ≈ 0.05) under the control scenario of unbiased even recording, including the *Naïve* model (figure 1; Appendix S2). The simple version of *ListLength* and *RepoitingRate* methods return significant results around twice as frequently as expected: this behaviour is fixed by adding a random effect for site identity (the *+Site* variants). The three methods that split the data into two time-periods (*RDC, Telfer* and *Frescalo_P*) are all conservative (α < 0.05): indeed RDC almost never rejected the null hypothesis across all parameter combinations (figure S1).

**Figure 1.**
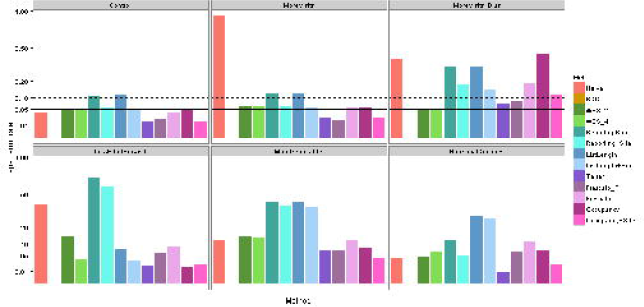
Type I error rates of all methods under all scenarios (note square root scale on y-axis). Results are shown for medium levels of recording intensity. The solid and dashed lines indicate α=0.05 and α=0.1 respectively.

All methods experience at least one combination of recording scenario and input parameters in which the Type 1 error rate is inflated by a factor of two compared with the *Control* (figure 1, table S1). Under three scenarios (*MoreVisits+Bias, MoreDetectable, NonFocalDeclines),* the failures become more acute as the quantity of data increases (figure S1), reflecting the fact that small datasets contain insufficient data to reject the null hypothesis.

As predicted, the *Naïve* model performs badly under virtually all departures from random sampling. Other methods are robust to growth in the number of visits (*MoreVisits*), i.e. the Type 1 error rate is close to that observed under the *Control.* The performance of several methods deteriorates markedly when in our spatial biased scenario (*MoreVisits+Bias),* notably *Frescalo_Y, RepoitingRate+Site, ListLength+Site* and both implementations of *Occupancy.*

When recording becomes progressively more incomplete (*LessEffortPerVisit*), the *ReportingRate+Site* and *WSS_2* both fail, reflecting the fact that it becomes increasingly less likely that the focal species will be recorded on an average visit. Increasing the threshold list length solves the problem (*WSS_4*), as predicted. Both implementations of *Frescalo* and *Occupancy* are robust to this form of bias, although the latter is conservative.

Changes in detectability (*MoreDetectable*) elevate Type 1 error rates in almost all methods. For *Occupancy, Frescalo_P* and *Telfer* the elevation is slight (α < 0.1 under all levels of recording intensity), but the failure is more extreme for *WSS* and *Frescalo_Y,* especially under high recording intensity (figure S1). *NonFocalDeclines* induce poor performance of *ListLength+Site* and *Frescalo_Y,* but only slight elevations for both implementations of *Frescalo_P* and *Occupancy.*

In summary, the *Naïve, RepoitingRate* and *ListLength* models (including *+Site* variants) all experience serious failures under a majority of biased recording scenarios and are therefore not robust.

Not surprisingly, power is strongly affected by overall sampling intensity, with a two-fold increase going from low to high intensity recording (figure 2). Power declines under most deviations from the *Control* (figure 3), but the relative power of each method is fairly consistent, with *Occupancy+Site* being most powerful, followed by the simple version of *Occupancy,* then *Frescalo_Y, Frescalo_P, Telfer, WSS_2, WSS_4* and finally *RDC* (which has virtually no power at all). The exceptions to this rule are *LessEffortPerVisit,* in which case *Frescalo* outperforms *Occupancy,* and *NonFocalDeclines,* in which *WSS* outperforms *Frescalo* (figure 3, figure S2).

**Figure 2:**
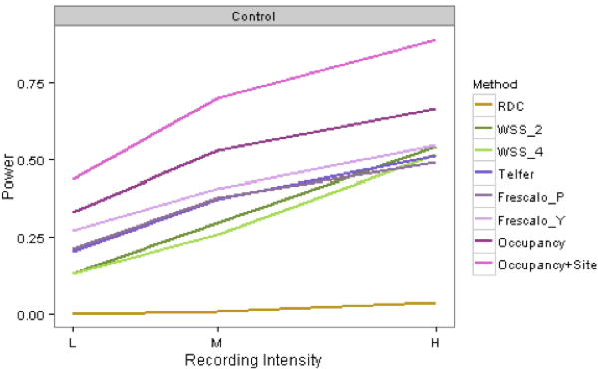
Power under the Control scenario plotted against recording intensity. Results are not shown for five methods that failed the test of validity.

**Figure 3:**
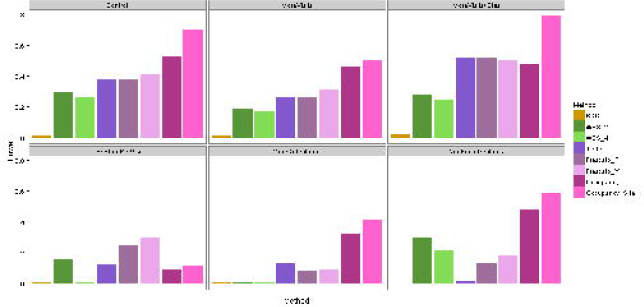
Power under medium recording intensity for all scenarios. Results are not shown for five methods that failed the test of validity.

## Discussion

Our simulations have provided a rigorous test of candidate methods for estimating trends in species’ distributions from opportunistic data. Many studies have emphasised the problem that opportunistic data were generated with uneven sampling effort over time (Botts et al., 2012; Maes et al., 2012; Prendergast et al., 1993), but we observe that most methods are robust to this (*MoreVisits* scenario). Other forms of variation in recorder activity present serious problems for many methods, yet are rarely discussed. We found that none of the methods is robust under all scenarios, but several perform well enough to be useful, and some general principles have emerged about how to apply then to real-world datasets.

We have clear evidence that simple methods easily fail under realistic scenarios of recording behaviour. The poor performance of the *Naïve* model is not unexpected, but the *ReportingRate* and *ListLength* (including *+Site* variants) both failed under a majority of scenarios (table 3). The simple versions of both methods failed even under the *Control* scenario of random sampling, since they treat visits as independent. Our findings draw into question the conclusions of studies that have used such methods (Breed et al., 2012; Szabo et al., 2011). The trend estimates from these methods are likely to be unreliable in any situation where the sampling variance of the focal species is high or uneven, including when the timescale is short, and when the study area is large and/or heterogeneous. The *RDC* method fails in a different way: it almost never rejects the null hypothesis (because few sites qualify as well-sampled) and always under-estimates the true trend (because data are aggregated into time periods). These features imply that published trends (e.g. Maes et al., 2012) are highly conservative. *Telfer’*s method, which is also relatively simplistic, performed consistently well but never better than *Frescalo_P,* which produces trends that are easier to interpret.

**Table 3:**
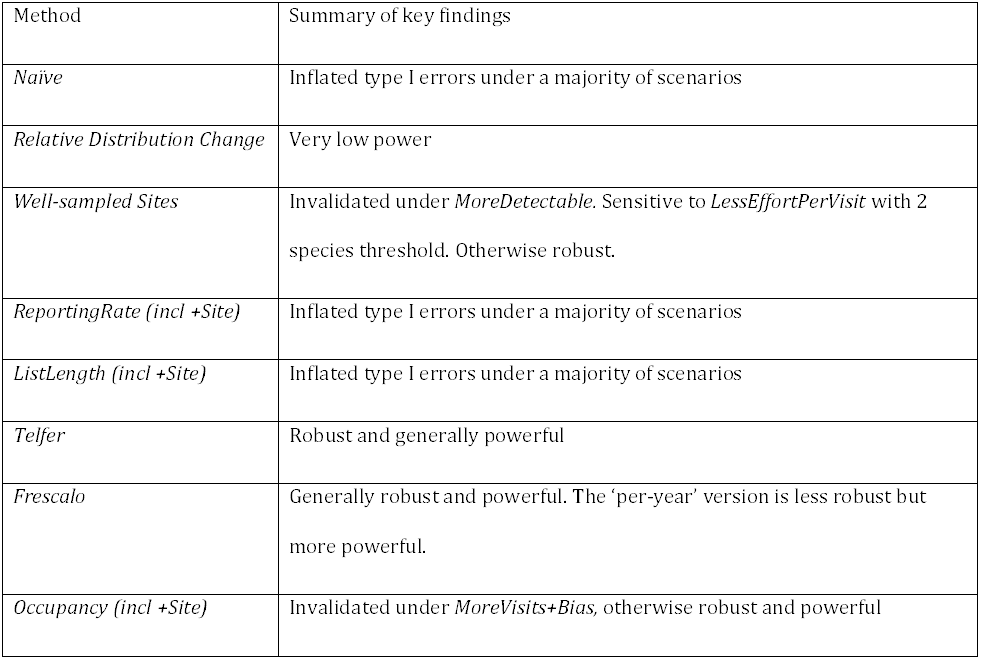
Summary of method performance across all tests

Previous studies have compared only simple methods (Botts et al., 2012), but our results show that complex methods outperform simple ones. In some cases, the reasons for this strong performance are clear: models with Site effects are more robust than those without; *Occupancy* is the most robust method under *More Detectable* because it explicitly models the detection process; *WSS_4* (but not *WSS_2*) is robust to *LessEffortPerVisit* because visits with low effort (defined here as L<4) are excluded. By contrast, we were surprised that *Frescalo_P* (although not *Frescalo_Y*) is reasonably robust to scenarios where the focal species undergoes separate treatment (*MoreVisits+Bias, MoreDetectable*). A deeper understanding of why methods fail, and why they perform well, would help develop methods that are robust to all forms of variation in recorder activity.

Until such universally robust methods become available, we must devise tests to determine the extent to which real datasets exhibit the specific forms of bias modelled here. It should be possible to diagnose whether the prevalence of short lists changes over time, or whether the spatial footprint of recording has shifted with respect to the focal species’ distribution. Changes in detectability are likely to be more challenging, because detection is a function of both the species’ ecology and the data collection process (Isaac et al., 2011; van Strien et al., 2013). In the absence of a single best method, we are encouraged that the three best performers (*WSS_4, Frescalo_P, Occupancy+Site*) have complementary strengths and weaknesses. Thus, one approach to trend estimation would be to draw inferences from an ensemble of methods (c.f. Thuiller et al., 2009). Our experience to date is that trends from different methods broadly agree (Isaac et al., 2013).

Overall, we feel that sophisticated methods such as *Frescalo* and *Occupancy,* which model the data collection process, have the greatest potential for delivering robust and timely trends from opportunistic data. Selection methods, including *WSS,* are ultimately limited by the assumption that simple thresholds can separate the signal from the noise, and by the loss of power that results from discarding data (at least 75% of site:year combinations in most simulations). However, selection criteria may still have a role in addressing specific forms of bias that are difficult to model. For example, excluding sites with few years of data (as employed by *WSS*) could be an effective solution to the problem of spatial bias in site selection that produced inflated type 1 errors for *Occupancy.*

Whilst *Frescalo_P* performed well in our simulations, we have a number of reservations about its usage. First, using the method requires the user to make a variety of choices, in addition to the number of time periods. The selection of benchmark species and neighbourhoods are defined by input parameters (Hill, 2012) which have considerable impact on the trend estimates that are produced (van Strien et al., unpublished data). Second, our simulations compared all methods at the same spatial scale, but the typical grain size for *Frescalo* is 100-fold larger (100 km^2^ *vs* 1 km^2^) than used by *WSS* (Roy et al., 2012) and *Occupancy* (van Strien et al., 2010, 2013), so the number of unique observations (and hence power) is also lower. This coarse-grained approach reflects both computational limitations (neighbourhoods are defined by a matrix of N x N, where N is number of sites), and the need to robustly estimate recording intensity for each site. However, *Frescalo* remains the most appropriate method for describing long-term change where the periods are well-defined (e.g. published atlases) and when information from individual visits is unavailable (Hill, 2012).

We modelled a suite of recording scenarios, but there is a gap between our idealised simulations and the reality of how opportunistic data are collected. Our four axes of variation in recorder activity conceal many specific departures from the central assumption that species are recorded as complete assemblages during site visits. This assumption is violated during targeted surveys, or where recorders make annual lists but submit records from individual visits: in this case species reported during early visits get omitted from lists made later in the year. At present we lack information about how the records were generated, such as whether all observations were reported. The growth of technology in wildlife recording, including smartphone apps, offers great potential to capture meta-data about sampling intensity (e.g. start and end times of the survey) with minimal input from the recorder. These data would go a long way to make inferences from opportunistic data more robust in future.

Our results add to a growing body of evidence that opportunistically-gathered data has enormous potential to make meaningful contributions in biodiversity science and policy-making (Schmeller et al., 2009; Tulloch et al., 2013). Some of the methods we tested here (e.g. *WSS, Occupancy*) can easily incorporate covariates, making them ideal for testing hypotheses about the drivers of biodiversity change (e.g. Roy et al., 2012). Our results provide an evidence base for producing quantitative trends from opportunistic data and a benchmark against which future methods can be compared.

## Acknowledgements

We are grateful to Gary Powney for constructive comments on a previous version of this manuscript, to Stuart Ball and Mark Hill for advice, and to Stephen Freeman, Colin Harrower and Thierry Onkelinx for technical advice. This work was funded by JNCC, NERC and the Welsh Government.

## Data Accessibility

All computer code to run the simulation and draw the figures is available at https://github.eom/BiologicalRecordsCentre/RangeChangeSims.

### Appendices

Appendix S1: Statistical description of the methods compared by simulation

Appendix S2: Detailed results of the simulation study

